# Population Genomics of Rapidly Invading Lionfish in the Caribbean Reveals Signals of Range Expansion in the Absence of Spatial Population Structure

**DOI:** 10.1101/291666

**Authors:** Eleanor K. Bors, Santiago Herrera, James A. Morris, Timothy M. Shank

## Abstract

Range expansions driven by global change and species invasions are likely to have significant genomic, evolutionary, and ecological implications. During range expansions, strong genetic drift characterized by repeated founder events can result in decreased genetic diversity with increased distance from the center of the historic range, or the point of invasion. The invasion of the Indo-Pacific lionfish, *Pterois volitans*, into waters off the U.S. East Coast, Gulf of Mexico, and Caribbean Sea provides a natural system to study rapid range expansion in an invasive marine fish with high dispersal capabilities. We report results from 12,759 loci sequenced by restriction enzyme associated DNA sequencing for nine *P. volitans* populations in the invaded range, including Florida and other Caribbean sites, as well as mitochondrial control region D-loop data. Analyses revealed low to no spatially explicit metapopulation genetic structure in the study area, which is partly consistent with previous finding of little structure within ocean basins, but partly divergent from reports of between-basin structure. Genetic diversity, however, was not homogeneous across all sampled sites. Patterns of genetic diversity correlate with invasion pathway. Observed heterozygosity, averaged across all loci within a population, decreases with distance from Florida while expected heterozygosity is mostly constant throughout sampled populations, indicating population genetic disequilibrium correlated with distance from the point of invasion. Using an F_ST_ outlier analysis and a Bayesian environmental correlation analysis, we identified 256 and 616 loci, respectively, that could be experiencing selection or genetic drift. Of these, 24 loci were shared between the two methods.

## INTRODUCTION

The distributions of species change over multiple temporal and spatial scales due to natural and human-driven processes, such as glacial retreat over interglacial periods (Hewitt 1999; 2000; Silva *et al.* 2014; Shum *et al.* 2015), global climate change (Parmesan & Yohe 2003; Perry 2005; Harley *et al.* 2006; Pinsky *et al.* 2013), local habitat alteration (Bradshaw *et al.* 2014), and non-native species invasions (Lowry *et al.* 2013). Range expansions, a common type of distributional shift, can result in decreased genetic diversity with increasing distance from the center of the original range (Excoffier *et al.* 2009), as has been observed in humans with distance from Africa (Ramachandran *et al.* 2005). Range expansion can also lead to evolution in life history traits (*e.g.,* Phillips *et al.* 2010). One genetic outcome of the spatial process of range expansion is allele surfing, (alternatively called “gene surfing” or “mutation surfing”).

Allele surfing is a process by which an otherwise rare allele or new mutation rises to high frequency or fixation near a moving range margin because of repeated founder events through space and time (Edmonds *et al.* 2004; Klopfstein 2005; Hallatschek & Nelson 2008; Peischl *et al.* 2013). The phenomenon of allele surfing is related to, but not exactly the same as, bottlenecks in diversity due to large founder events. Allele surfing can vary in strength, leaving either strong or subtle gradients in allele frequencies, and could potentially contribute to population genomic patterns during range expansion.

Invasive species are frequently studied in evolutionary biology as “natural experiments” or models to investigate the dynamics of invasion as well as adaptation to new environments (Barrett 2015). Being able to predict the evolutionary dynamics of range expansion during invasion may be important for managing invasions and anticipating impacts of climate-driven range shifts. Here, we use the invasion of the Indo-Pacific lionfish, *Pterois volitans* [Linnaeus, 1758] as a model for rapid range expansion on a decadal time scale in a marine species with high dispersal capabilities.

The invasion of *Pterois volitans* and *Pterois miles* [Bennett, 1828] in the Western Atlantic and Caribbean Sea is unprecedented in both rate of geographic spread and ecological damage (Albins & Hixon, 2011; Albins 2015; Hixon *et al.* 2016). First reported off Dania, Florida in 1985, the lionfish invasion in the western Atlantic likely originated in southern Florida, and has been characterized by a long incubation period and an immense post-establishment expansion (Morris & Akins 2009). In the late 1990s and early 2000s, lionfish began expanding northward; in 2004, they spread to the Bahamas; and in the years since, they have invaded the Caribbean Sea and the Gulf of Mexico (Schofield 2009; Schofield 2010, Ferreira *et al.* 2015) (summarized in Figure 1). *Pterois volitans* is the most common species in the invasion, with *P. miles* mostly restricted to the northern part of the invaded range (Freshwater *et al.* 2009). While there has been speculation that *P. volitans* and *P. miles* could hybridize in the invaded range, Wilcox *et al.* (2017) has recently presented phylogenetic evidence that the lineage we know as *P. volitans* in the invaded range may in fact represent a hybrid lineage between *P. miles* and another Pacific Ocean species of *Pterois.* The present study focuses on the *P. volitans* lineage as it has historically been defined, and will use that species name here, while recognizing that there may be unresolved phylogenetic treatment of *Pterois*.

**Figure 1.**
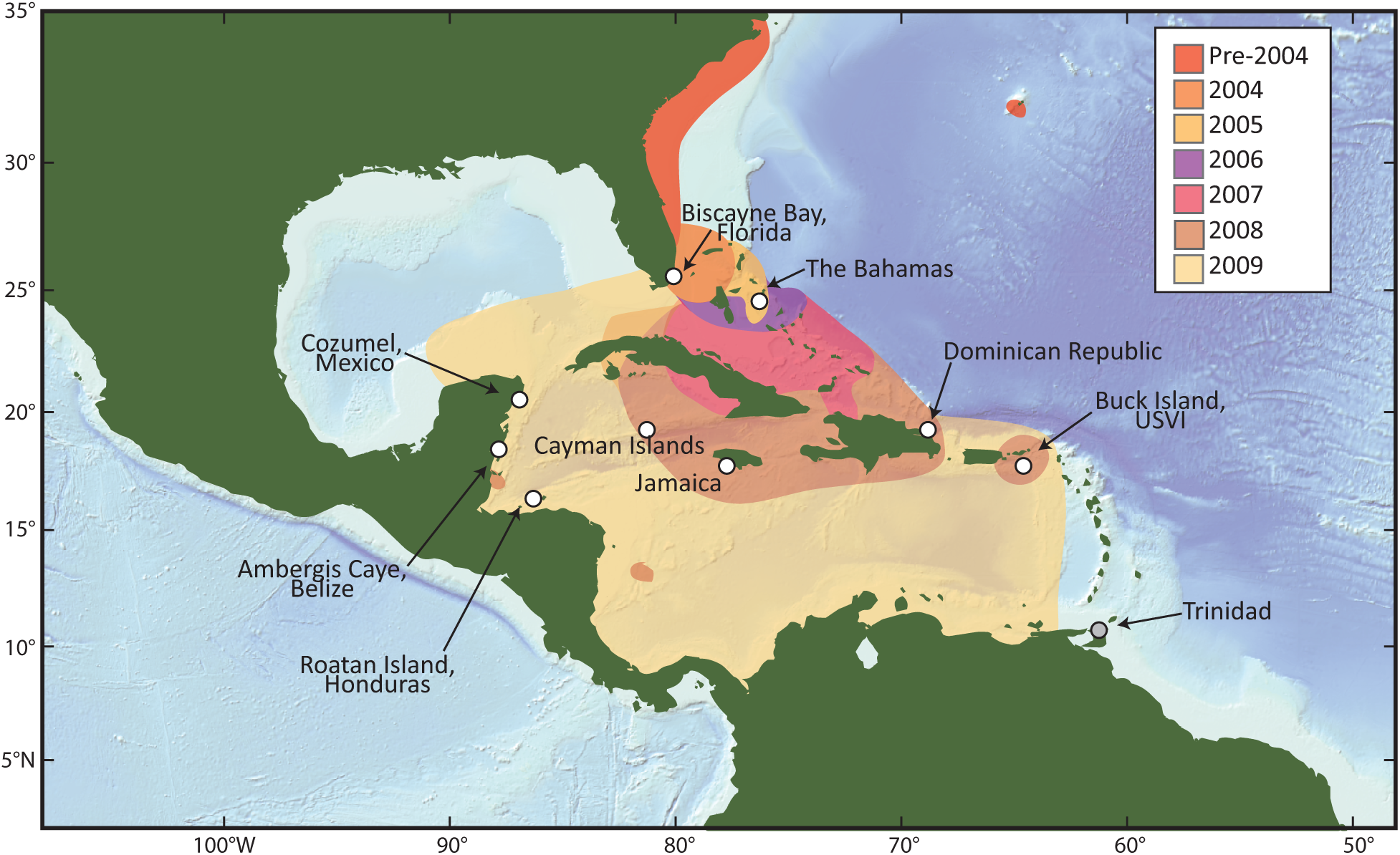
Map of the study region showing the nine sampling locations used for RAD-sequencing (samples from Trinidad, shown here with a gray circle, were used in mitochondrial analyses). Colored contours on the map indicate the extent of the invasion in the years from 2004-2009, by which point all of the nine sites had been invaded (see legend for dates).

For lionfish in the invaded range, recent genetic studies have focused on mitochondrial sequencing to describe population genetic connectivity and population structure. To date, studies have identified just nine haplotypes of mitochondrial D-loop in the invaded range but have not traced these directly back to a specific source in the native range, where genetic diversity is much greater (Freshwater *et al.* 2009; Betancur-R *et al.* 2011; Butterfield *et al.* 2015; Johnson *et al.* 2016). While north-to-south (*i.e.,* Western Atlantic to Caribbean) population differentiation has been reported in the invaded range, overall, a lack of metapopulation genetic structure has been reported within oceanic basins (Freshwater *et al.* 2009; Betancur-R *et al.* 2011; Toledo-Hernández *et al.* 2014; Butterfield *et al.* 2015; Johnson *et al.,* 2016), with some local population structure reported in Puerto Rico (Toledo-Hernández *et al.* 2014). Most recently, evidence for a bottle neck between the Caribbean and the Gulf of Mexico populations has been reported, evidenced by the presence of only three of the four Caribbean D-loop haplotypes in Gulf of Mexico populations (Johnson *et al.* 2016). Johnson *et al.* (2016) provides evidence for two founder events when lionfish entered each new basin, potentially caused by gene flow restrictions related to oceanographic barriers. These course-scale patterns are congruent with large-scale barriers to dispersal between oceanic basins that could have restricted gene flow into new areas as lionfish invaded. Gradients in major allele frequency in populations across the lionfish’s invasion pathway are generally beyond the scope of single locus mitochondrial studies but can be achieved by sequencing sites located throughout the genome.

The use of next generation sequencing (NGS) and other emerging genomic tools to provide novel insights into the evolutionary repercussions of range expansions and invasion dynamics is now widely recognized as the frontier in invasion genetics research, promising a synergy between previously intractable questions and burgeoning technologies (Kirk *et al.* 2013; Chown *et al.* 2014; Rius *et al.* 2015; Barrett 2015; Bock *et al.* 2015). Recent examples have demonstrated the power of NGS datasets to identify genomic regions undergoing neutral evolution and regions subject to natural selection with potential adaptive roles during an invasion (White *et al.* 2013; Tepolt & Palumbi 2015). In one such study, White *et al.* (2013) found evidence of both genetic drift and natural selection in populations of an invasive Bank Vole in Ireland, including patterns of decreased genetic diversity towards the moving range edge.

This study presents genome-wide single nucleotide polymorphism (SNP) data for the invasive lionfish collected throughout the Caribbean Sea, using 12,759 loci across nine populations. These data are analyzed from a range expansion perspective, identifying changes in genetic diversity with distance from the point of invasion. We test the hypothesis that range expansion of lionfish is characterized by repeated founder effects at the moving range edge leading to decreased genetic diversity (major allele frequency, allelic richness, and heterozygosity) with increased distance from the point of introduction in Southeastern Florida. We predicted that signals of allele surfing would be detectable in the SNP data, in line with the theoretical predictions outlined above. Counter to these predictions, patterns of decreased diversity in the form of decreased major allele frequency or allelic richness were not observed in the data. However, decreases in average observed heterozygosity were observed, indicating higher levels of non-equilibrium population genetic dynamics near the range edge, with more central populations tending towards an equilibrium state. This may occur as populations become more established and the invasion front propagates forward. We also identify outlier loci in both Bayesian and F_ST_ analyses that may be under drift or under selection, potentially playing adaptive roles in certain lionfish populations.

## Methods

### Sample collection

Pterois volitans individuals were collected from nine Caribbean sites for genomic analysis (Figure 1). Additional individuals were collected from Trinidad and included in mitochondrial analysis but not in RAD-seq analysis. *Pterois volitans* specimens from Biscayne Bay, Florida, were collected by SCUBA divers from the U.S. National Park Service in August and September of 2013 as part of ongoing collection programs. Fin clips were subsampled from each fish and stored in ethanol in a −20°C freezer. Similarly, samples from the U.S. Virgin Islands were collected from Buck Island by divers from the University of the Virgin Islands between May of 2013 and February of 2014 and fin clips were subsampled. Samples from The Bahamas, the Dominican Republic, Jamaica, the Cayman Islands, Cozumel (Mexico), Belize, Honduras, and Trinidad were collected by divers throughout 2013 and tissue subsamples were archived in the U.S. National Oceanic and Atmospheric Administration (NOAA) Beaufort Laboratory, Beaufort, North Carolina. Samples from Trinidad were only used in mitochondrial analyses due to DNA quality requirements for genomic analyses. All other sites were used for both mitochondrial and RAD-sequencing analysis. Sections of muscle tissue from archived filets were subsampled at NOAA. Fish were identified to species when possible through meristics (*i.e.,* morphological traits) at the collection site and later confirmed through molecular barcoding. If provided by collectors, latitude and longitude, depth, date of collection, sex, and standard or total length for each sample are given in SI-1. The latitude and longitude of the most common collection site per country were used in subsequent spatial analyses (SI-1). Tissue samples were shipped to the Woods Hole Oceanographic Institution in ethanol or frozen and then stored at −80°C until genomic DNA extraction.

To estimate the age of each sampled individual, and therefore the likely time of recruitment of the individuals used in this study, we calculated age from total length using a von Bertalanffy growth curve (Barbour *et al.* 2011). For samples that lacked a standard length measurement but had a total length measurement, we utilized a conversion function to estimate standard length (Fogg *et al.* 2013). Distributions of estimated fish age and recruitment year are presented in SI-1.

### DNA extraction, and mitochondrial DNA PCR, sequencing, and analysis

Genomic DNA (gDNA) was extracted from muscle or fin clip tissue using a CTAB and proteinase K digest, a phenol-chloroform purification, and an ethanol precipitation as described in Herrera *et al.* (2015b). Genomic DNA was stored in AE buffer from a QIAGEN DNeasy Blood and Tissue Extraction Kit (Qiagen GmbH, Germany) at 4° C or −20° C until gene amplification and sequencing.

Polymerase chain reactions (PCRs) were performed targeting the mitochondrial control region D-loop with primers LionfishA-H (5’-CCATCTTAACATCTTCAG TG-3’) and LionfishB-L (5’-CATATCAATATGATCTCAGTAC-3’) (Freshwater *et al.* 2009). The thermocycler temperature profile consisted of 95° denaturing step for 3.5 minutes, then 30 cycles of 95° for 30 seconds, 51° for 45 seconds, 72° for 45 seconds, followed by a final extension step at 72° for 5 minutes. PCR reactions were purified using a QIAGEN PCR Purification Kit (Qiagen GmbH, Germany) and were sequenced in both directions using Sanger sequencing at Eurofins Operon Genomics (Eurofins MWG Operon LLC, Louisville, KY, USA). Sequences were edited and aligned using *Geneious 8.1.5* (http://www.geneious.com, Kearse *et al.* 2012) and were compared to the previously published haplotypes. Mitochondrial sequence data were generated for a total of 217 individual *P. volitans* samples (23 from Florida, 17 from The Bahamas, 16 from the Dominican Republic, 25 from Jamaica, 15 from the Cayman Islands, 24 from Mexico, 18 from Belize, 24 from Honduras, 23 from the U.S. Virgin Islands, and 32 from Trinidad). Genome-wide single nucleotide polymorphism (SNP) data were generated for a subset of 120 samples (Table 1; Table SI I.2).

**Table 1.** Population genomic summary statistics averaged over all loci and by population, as generated by the *Stacks*_*populations* program. N = number of individuals sequenced, N (Stacks) = average number of individuals used across all sampled loci, Private = number of private alleles in the population, P = major allele frequency (average), Obs Het = observed heterozygosity, Exp Het = expected heterozygosity, Pi = nucleotide diversity, F_IS_ = inbreeding coefficient.

| Pop ID | N | N (Stacks) | Private | P | Obs Het | Exp Het | Pi | F<sub>IS</sub> |
| --- | --- | --- | --- | --- | --- | --- | --- | --- |
| *FLO* | 11 | 10.4786 | 421 | 0.9053 | 0.1177 | 0.1334 | 0.1402 | 0.0643 |
| *BAH* | 9 | 8.6021 | 620 | 0.9024 | 0.0997 | 0.139 | 0.1476 | 0.1304 |
| *CAY* | 11 | 10.0495 | 478 | 0.909 | 0.0718 | 0.1283 | 0.1351 | 0.1786 |
| *JAM* | 14 | 13.0478 | 746 | 0.9077 | 0.0873 | 0.1316 | 0.1369 | 0.1498 |
| *DOM* | 16 | 14.6043 | 656 | 0.907 | 0.0802 | 0.1325 | 0.1373 | 0.1779 |
| *MEX* | 7 | 6.5103 | 381 | 0.9096 | 0.0772 | 0.1261 | 0.1367 | 0.1474 |
| *BEL* | 20 | 17.7624 | 839 | 0.9057 | 0.0678 | 0.1338 | 0.1377 | 0.2272 |
| *HON* | 15 | 14.6673 | 781 | 0.9056 | 0.0882 | 0.1348 | 0.1397 | 0.1564 |
| *USV* | 16 | 13.7671 | 857 | 0.9069 | 0.0788 | 0.1329 | 0.1379 | 0.1828 |

### Restriction enzyme associated DNA sequencing

Restriction enzyme associated DNA sequencing (RAD-seq) library preparation using the *SbfI* restriction enzyme (restriction site: 5’-CCTGCAGG-3’) was carried out on concentration-normalized gDNA by Floragenex Inc. (Eugene, OR, USA) in identical fashion to several other recent RAD-seq studies (Reitzel *et al.* 2013; Herrera *et al.* 2015b). Genomic DNA was digested with the *SbfI* restriction enzyme, yielding fragments of many different lengths. Barcode tags (specific to each individual) 10 basepairs (bp) in length and an Illumina adaptor, were ligated onto the sticky end of the cut site. Samples were then pooled, sheared, and size selected for optimal Illumina sequencing (Illumina Inc., San Diego, CA, USA). A subset of samples were prepared for paired-end Illumina sequencing (Illumina Inc., San Diego, CA, USA) following the library prep protocol described in Baird *et al.* (2008), in order to generate longer sequencing assemblies for future analyses as well as provide possible comparisons of methods. For the preparation of the paired-end sample library, a second adaptor was ligated to the second end of the read. All libraries were then enriched through PCR and sequenced by 96-multiplex in a single lane of an Illumina Hi-Seq 2000 sequencer (one lane for the single end sequencing, one for the paired end). For the samples sequenced in a paired-end Illumina run, each sample was loaded twice to achieve a standard coverage (*i.e.,* for one individual, two libraries were generated from two aliquots of gDNA with two barcodes).

### RAD-seq data processing and population genomic analyses

Using the *process_radtags* program in *Stacks* v1.19, raw Illumina, reads were filtered for quality with a minimum phred score of 10 in a sliding window of 15% read length (default settings) and sorted by individual-specific barcode. Reads were truncated to 90 bp, including the 6 bp restriction site. For the data generated with paired-end sequencing, only the first read was used. Putative loci were generated using the *denovo_map.pl* pipeline in *Stacks* v1.35 (Catchen *et al*. 2011) (references to *Stacks* from this point forward will all be to this version). We used a *stack-depth parameter* (*-m*) of 3, such that 3 reads were required to generate a stack (*i.e.,* a locus); a *within-individual distance parameter* (*-M*) of 3, allowing for three SNP differences in a read; and a *between-individual distance parameter* (-*n*) of 3, allowing for three fixed differences between individuals to build a locus in the catalog. In initial exploratory analyses, altering the values of the *within-individual* and *between-individual* parameters did not significantly impact the number or identity of downstream loci called by *Stacks* (not reported).

Population summary statistics (allele frequencies, observed and expected heterozygosities, π, and F_IS_) were calculated by the *populations* program in *Stacks*, using loci found in eight of the nine populations and in at least 80% of individuals per population (command flags *-p* 8, *-r* 0.8). Information on the effect of changing the *-p* and *-r* flags is available in SI-2. For each RAD-tag, only one SNP was used from 90 bp sequence using the program flag *–write_random_snp* (if there were two or more SNPs in the sequence, *Stacks* would randomly choose one to analyze). Heterozygosity (observed and expected) values were also calculated in the R Package PopGenKit (R Core Team, 2016; https://cran.r-project.org/web/packages/PopGenKit/index.html) to provide secondary validations of reported values. Allelic richness was calculated using PopGenKit.

Three methods were used to describe the genetic structure of lionfish populations in the study area: principal component analysis (PCA), a *STRUCTURE* analysis, and F_ST_ calculations. The *smartpca* program in *EIGENSOFT* (Price *et al.* 2006) was used to perform a PCA of genetic diversity. Custom iPython notebooks used to convert *Stacks* PLINK output files into *EIGENSOFT* input files, and for the visualization of the PCA are available at the author’s GitHub (https://github.com/ekbors/lionfish_pop_gen_scripts). *Smartpca* was run with four iterations of outlier removal (‘*numoutlieriter’* = 4) with otherwise default parameters. In addition to the PCA analysis, *fastSTRUCTURE* (Pritchard *et al.* 2000; Hubisz *et al.* 2009; Raj *et al.* 2013) was run with the number of genetic lineages (the value of *k*) set to values between one and ten to assess genetic structure through a hierarchical analysis, and the program *chooseK.py* was run to select the value of *k* most consistent with the program’s spatial structure model. F_ST_ values were calculated by the *populations* program in *Stacks* using a p-value cutoff of 0.05 and a Bonferroni correction (using the *‘bonferroni_gen’* flag in the *populations* program). In addition to these analyses, frequency spectra of the major alleles and of F_IS_ reported by *Stacks* were plotted in iPython. F_IS_ is calculated as 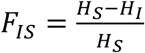 where H_S_ is the heterozygosity in the subpopulation and H_I_ is the heterozygosity of the individual. The output from *Stacks* reports F_IS_ values of zero when the H_S_ is equal to zero (*p* = 1), but in these cases, the numerical value of F_IS_ is actually undefined. In order to remove these values, only F_IS_ values that were calculated when H_EXP_ > 0 were used.

Genetic diversity summary statistics were regressed against distance from the southern Florida collection site of Biscayne Bay using the *stats* package from *Scipy*, a collection of open source software for scientific computing (https://scipy.org). The least cost distance dispersal trajectories used in these regressions were calculated using the ‘gdistance’ package in R with a bathymetric constraint from ETOPO1 (van Etten, 2015; R Core Team, 2016) with an additional requirement that pathways to sites to the west of Cuba first went around the east side of Cuba, a reasonable alteration considering the direction and strength of the Florida Current, as well as existing literature about the difficulty of dispersal of lionfish across that current (Johnston & Purkis 2015). Other methods of measuring distance were explored, including Euclidian distance and a non-modified least-cost ocean distances (that did not require pathways to go around Cuba) that result in slightly different regressions but ultimately the same conclusions (SI-3).

In addition to the described approaches of regressing genetic diversity measurements with distance from Florida, we also implemented range-expansion specific analyses (Peter & Slatkin 2013). Using an R package developed by Peter and Slatkin (2013), we calculated *psi*, or the “directionality index,” which measures asymmetries in allele frequency data to evaluate the likely direction of expansion in a set of populations and the relative distance of a site to the center of the range (no prior definition of the origin of expansion is needed).

### Blast2GO and locus identification

To annotate the RAD loci and infer possible links to gene function, we aligned the sequences to the non-redundant sequence database (restricted to teleost bony fishes) of NCBI using the BLASTx (Basic Local Alignment Search Tool) program as implemented in *Blast2GO* v2.5.1 (Conesa *et al.* 2005). We used an e-value threshold of 1×10^−3^, a word size of three and a HSP length cutoff of 33. BLAST results were used to map Gene Ontology (GO) and annotate RAD loci.

### Genome size estimation

To predict the size of the *P. volitans* genome based on the observed number of restriction sites (*i.e.,* half the number of observed RAD loci), we used the linear model and parameter estimates for the *SbfI* enzyme described by Herrera *et al.* (2015a) as implemented in the program *PredRAD* (Herrera *et al.*, 2015a). To generate a range for the number of restriction cut sites for *SbfI*, we ran the *Stacks* pipeline and *populations* program with several different permutations of parameters (SI-2) and then used a range of the number of total RAD loci generated by the different program runs.

### Locus-specific diversity analyses

Custom scripts were developed to identify groups of loci in the data with unique diversity patterns (https://github.com/ekbors/lionfish_pop_gen_scripts). Loci were identified for which (1) the major allele switched to the minor allele in at least one of the nine populations (*i.e.*, “*p*” of the Hardy-Weinberg equation drops below 0.5), or (2) the difference between the maximum and minimum value of the overall major allele among the populations exceeded a defined value (measured at values of 0.5, 0.6, 0.7. 0.8, and 0.9). Loci identified by these filtering techniques were used in analyses of site frequency spectra and F_IS_ to determine if specific loci were driving and/or breaking patterns in the dataset, meaning that the forces driving those loci might be dominating the overall population data.

### LOSITAN and BayEnv outlier analyses

To detect genomic outliers potentially under selection or strong genetic drift driven by expansion (which will yield similar diversity patterns), we used two analysis programs. *LOSITAN* (Antao *et al.* 2008) utilized data-wide F_ST_ values to identify loci that were outliers in their F_ST_ values. We ran 1,000,000 simulations in *LOSITAN* for all nine populations with the options for “Forced mean F_ST_” and “Neutral F_ST_” selected. The false detection rate was set to 0.01 and a correction was implemented by the program.

The second program used for outlier analysis was *BayEnv 2.0* (Coop *et al.* 2010; Günther & Coop 2013), a program based on a Bayesian analysis that first develops a covariant matrix as a null model and then generates a linear model of relationship between diversity and an environmental factor. We used the calculated ocean distance from Florida as an environmental gradient against which to test patterns of diversity in the data. *BayEnv* controls for underlying population structure by generating a Bayes Factor for each locus indicating its relative goodness-of-fit to the linear model related to the environmental gradient. To interpret Bayes Factors, loci were binned in decimal intervals (randomly choosing *p* or *q* for each locus). Within each bin, each locus was ranked by Bayes Factor and that rank was divided by the number of loci in the bin. This created the empirical distribution from which loci in the top 5% and 1% of Bayes Factor values were identified, as described in Coop *et al.* (2010) and Hancock *et al.* (2010).

Traditionally, these analyses are used to identify regions of the genome under selection. However, signals of allele surfing and strong genetic drift in the case of a range expansion could lead to allele frequency patterns correlating with distance or with expansion in ways that resemble the patterns of selection. As mentioned in the Introduction, these signals could vary in strength. Therefore, in some cases, the loci showing correlation to the gradient of distance may just as likely be the result of drift as selection (*e.g.,* White *et al.* 2013).

## RESULTS

### No evidence of recent re-introductions in mitochondrial data

Mitochondrial haplotypes consisting of 679 bp of the mitochondrial control D-loop region were sequenced for 217 samples. Only 5 of the 9 known haplotypes previously described were identified in these samples (Freshwater *et al.* 2009; Betancur-R *et al.* 2011; Butterfield *et al.* 2015). These haplotypes correspond to previously-named haplotypes H01, H02, H03, H04, and H06. Mitochondrial data do not indicate any new introductions of genetic material since the first publication of mitochondrial population genetic data in 2009 (Freshwater *et al.* 2009). Also in line with previous studies, distributional patterns and haplotype relationships largely corresponded to those described in Butterfield *et al.,* 2015. For most locations, only 2 or 3 haplotypes were present in the tested sample, but all 5 haplotypes were found in the Bahamas samples. For a complete summary of the mitochondrial results, see SI-4.

### High-quality RAD-seq and single nucleotide polymorphism data

Processing of raw Illumina data by the program *process_radtags* in *Stacks* resulted in the removal of less than 1% of the data due to poor sequencing quality, about 20% of the data due to ambiguity in the restriction site, and between 9% and 16% of reads due to ambiguous barcodes (inability to attribute a sequencing read to an individual). The number of reads removed varied slightly by sequencing type (single end *vs.* paired end) and by population (SI-2). The mean depth of reads for each individual, averaged over loci, was 24.5 reads and the average of the standard deviations for each individual was 28.3. More in-depth information on the depth of coverage is provided in the supplemental information (SI-2). The *Cstacks* program in *Stacks* generated a catalog of 1,376,469 putative loci, 12,759 of which were used by the *populations* program and in all subsequent analyses. The overall patterns of genetic diversity and genetic structure were not altered significantly in different parameter runs of *Stacks*. When more loci were included in analyses, heterozygosity increased overall—trends held the same shape but shifted upwards. This filtering-diversity relationship is consistent with what is generally known about RAD-sequencing approaches specifically under-reporting diversity (Arnold *et al.* 2013) and being more conservative in the *populations* filtering for loci.

### RAD-Locus Identification

Blast2GO queries against all existing fish genome databases resulted in matches for 2,766 of the 12,759 loci (21.7%). In most cases, two RAD-tag sequences matched to a BLAST result, which is consistent with having two “loci” sequenced in each direction away from the restriction site. These results could be used in concert with future draft and scaffold assemblies of the lionfish genome to confirm identity and location or RAD loci.

### Genome size estimation and utilization

The number of RAD loci identified in multiple populations ranged from 9,502 to 48,079 with the majority of values between 30,000 and 50,000 (data are estimates from one catalog of loci generated by *denovomap.pl*, reviewed in SI-2). Given that sequencing in both directions from a cut site leads to two RAD “loci” at each cut site, we generated estimates for genome size for 15,000, 20,000, and 25,000 cut sites, representing the majority of putative values for cut sites (SI-5, Table SI.V.1). Estimates ranged from 370,725,631 bp to 680,784.288 bp. Considering these results, the 12,759 loci used in this study represent between 0.17% and 0.31% of the total lionfish genome.

### Genomic diversity correlates with the invasion pathway

Observed heterozygosity decreased linearly with distance from Florida (Figure 2A; Table 1) even though both allelic richness (average number of alleles per locus) and expected heterozygosity (calculated by *Stacks* as *2pq* from the Hardy Weinberg equation) remained steady throughout the sampled range (Figure 2B, 2C). The difference between the expected and observed heterozygosity—a measure of deviation from Hardy Weinberg equilibrium—increased with distance from Florida. All methods of measuring distance resulted in similar regressions for observed heterozygosity (SI-3). These range-expansion patterns were observed despite a notable lack of spatial metapopulation genetic structure.

**Figure 2.**
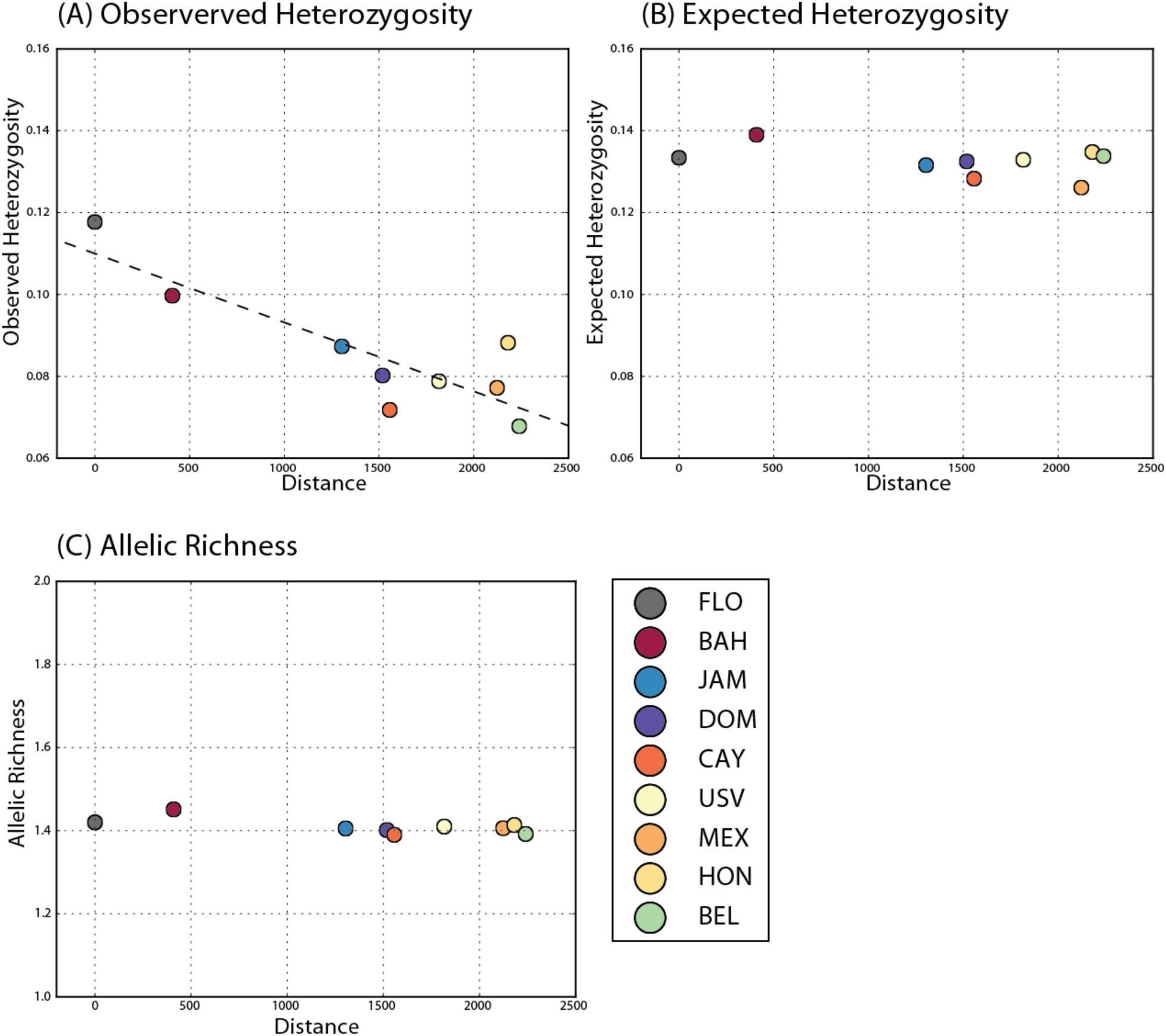
Summary statistics plotted against the “modified” ocean distance, measured from Florida. (A) Observed heterozygosity (R^2^ = 0.744, p-value = 0.003), (B) Expected heterozygosity (no significant regression), and (C) Allelic richness (no significant regression).

In general, site frequency spectra (SFS) distributions for each population were similar to each other (SI-6, Figure SI.VI.1), with some subtle variation. Many evolutionary and population processes can affect the shape of a SFS distribution and it is difficult to discern what could be driving such subtle differences (*e.g.,* Eldon *et al.* 2015). F_IS_ distributions in Florida and The Bahamas were closer to an equilibrium expectation of zero than F_IS_ distributions from populations closer to the moving range edge, which showed a thicker tail in the distribution skewing towards 1 (*e.g.*, the Cayman Islands, and Mexico) (Figure 3).

**Figure 3.**
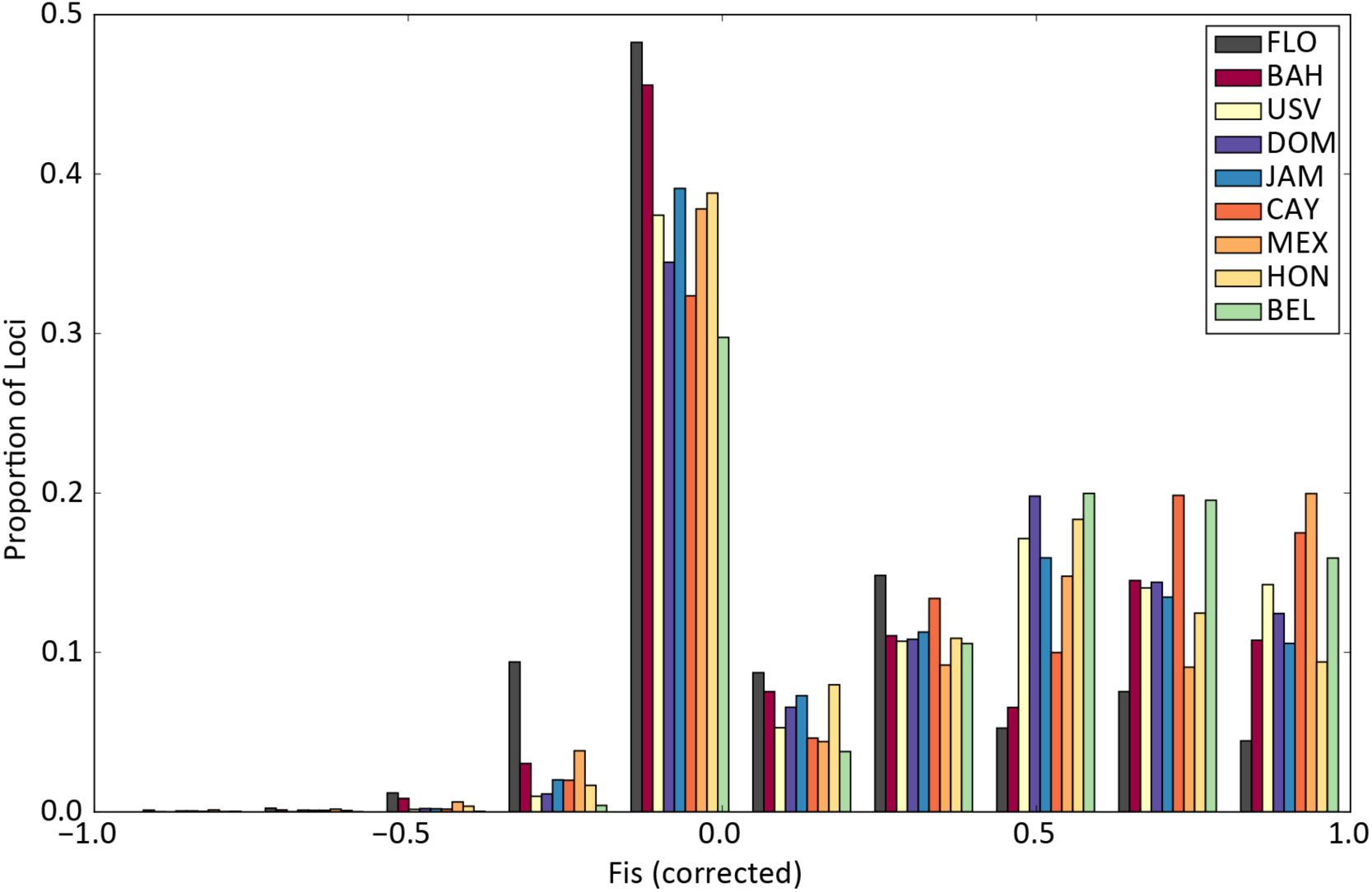
F_IS_ distributions showing the proportion of loci with F_IS_ values within each bin (number of bins = 10) for values between −1 and 1. Values of 0 reported by *Stacks* for loci for which the expected heterozygosity was 0 were removed from the data as described in the Methods.

### Lack of Spatially explicit metapopulation genetic structure

There was no obvious spatial metapopulation genetic structuring among the nine populations in the study region. Principal component analysis (Figure 4) revealed no clustering of defined populations with the first, second, and third components (*e.g.*, eigenvectors) accounting for 11.03%, 10.44%, and 10.30% respectively of the variation in the dataset. In order to determine the most likely number of genetic lineages (the value of *k*), or subpopulations, the *chooseK.py* program from *fastSTRUCTURE* was run for values of *k* between 1 and 10. The value of *k* that maximized marginal likelihood and that best explained the structure in the data (two different program metrics for assessing the appropriate value of *k*) was 1, indicating that the *fastSTRUCTURE* analysis fit the data best with just one genetic lineage. After a Bonferroni correction, many pairwise F_ST_ values calculated by *Stacks* were not statistically different from zero. For those that were, F_ST_ values showed very slight genetic differentiation among populations with significant values only for 5 pairings: Bahamas-Belize = 6.91 × 10^−5^; Caymans-Mexico = 1.1 × 10 ^−4^; Jamaica-Dominican Republic = 6.10 × 10 ^−5^; Jamaica-Honduras = 1.2 × 10 ^−4^; Dominican Republic-Honduras = 1.1 × 10 ^−^4. F_ST_ values, therefore, do not reveal genetic differentiation of populations closer to the edge from those at the center of the range.

**Figure 4.**
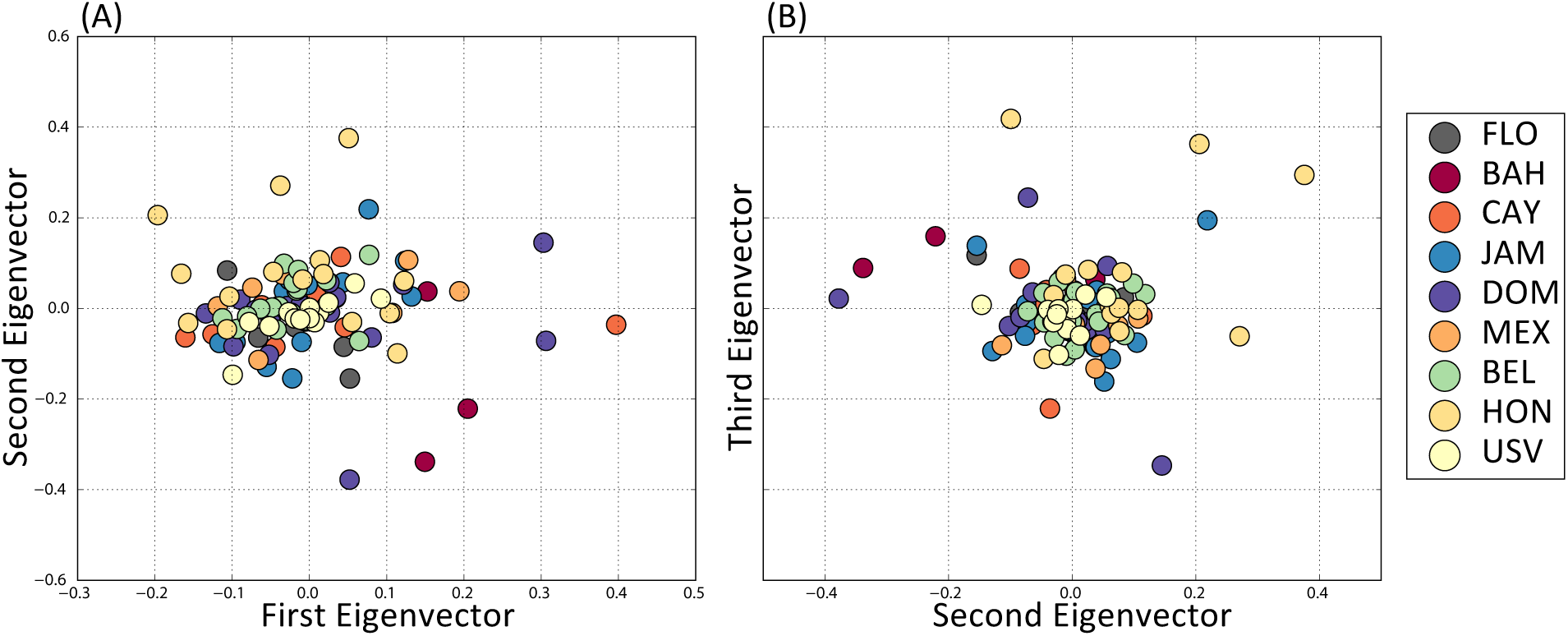
PCA generated by *smartpca* in *EIGENSOFT* here shown for the run with outliers removed.

The directionality index indicates another possible concept of distance from the point of invasion based on asymmetries of allele frequencies (Figure 5). The ordering of the index from lowest to highest indicates the “distance” in terms of the expansion from the center of the range. These data are ranked in the following order: Florida, Honduras, the Cayman Islands, U.S. Virgin Islands, Jamaica, The Bahamas, Mexico, Belize, and the Dominican Republic. This order of distance, or invasion directionality is different from an expectation based solely on geographic proximity. Specifically, the results indicate that the Dominican Republic is more isolated from the center of the invasion than all other sites, and that Honduras is much more connected to the core of the range even though it is geographically distant.

**Figure 5.**
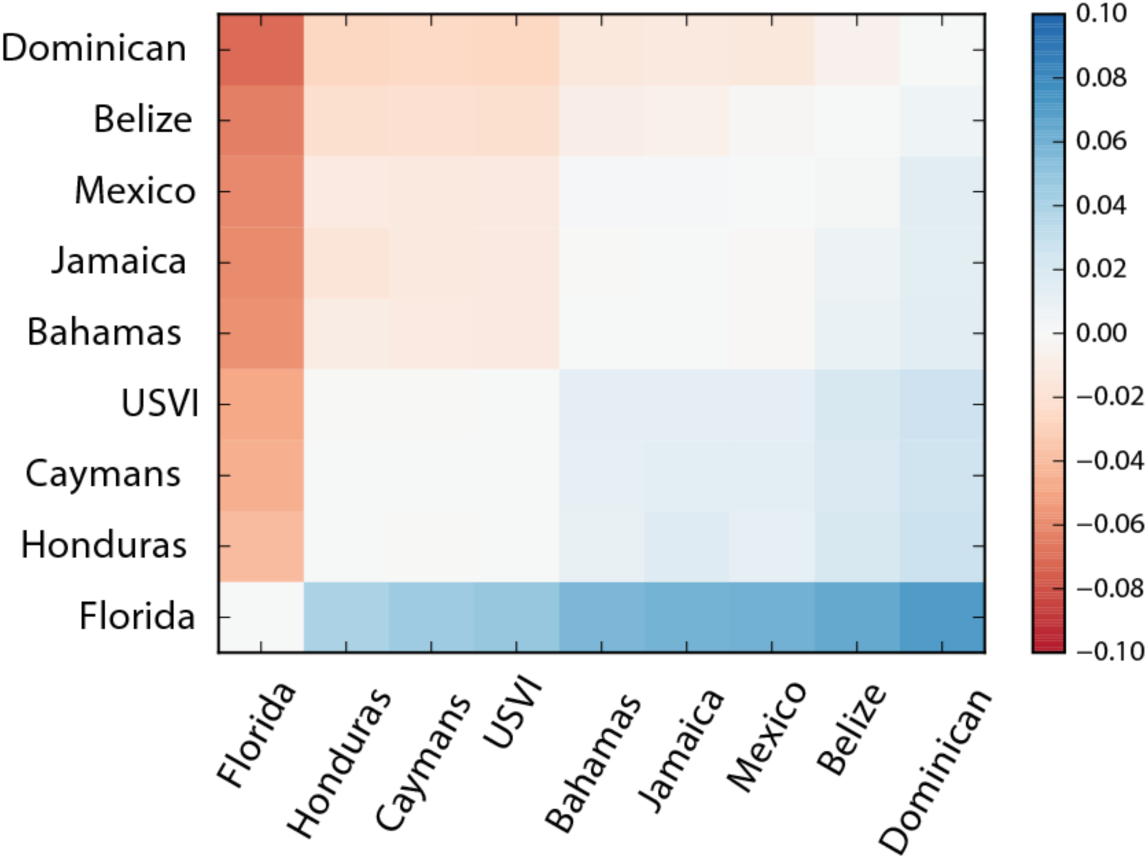
Directionality index heatmap. The directionality index, *psi,* measures asymmetries in allele frequencies. Here, values of *psi* have been arranged from lowest to highest—intended to parallel the ordering of sites from the closest to the origin of expansion to the furthest from the origin of expansion.

### Locus-specific patterns of spatial genetic diversity

There were 1,207 loci for which the value of *p*, or the major allele, defined as the allele most frequent across all the 120 samples, dropped below 0.5 in at least one population, meaning that for those loci, the major allele overall became the minor allele locally (called “Flip-Flop loci” here; Table 2). There were 290 loci with a difference in the minimum and maximum allele frequency of at least 0.5, 55 with a difference of at least 0.6, 3 with a difference of at least 0.7, and 1 with a difference of at least 0.8. There were no loci with a minimum-maximum difference of 0.9 or greater. Of the loci that switched from major allele overall to minor allele in at least one population, 243 were also present in the 0.5 difference list. Therefore, 964 of the loci that switched between being a major and minor allele never had a maximum difference that exceeded 0.5. These loci are likely oscillating around a frequency of 0.5, not demonstrating dramatic changes throughout the invaded range. Such loci are sometimes attributed to balancing selection. The 243 loci with larger differences between their minimum and maximum values, however, could be driven by specific forces such as drift and directional selection.

**Table 2.** Comparisons of the loci identified as outliers by the two outlier analyses and loci identified through different filtering methods through custom analysis presented in this paper.

| Overlap of BayEnv top 5% with other filtering methods (total #loci = 616) |  |  |  |  |  |
| --- | --- | --- | --- | --- | --- |
|  | Lositan | Flip-Flop | 0.5 diff | 0.6 diff | 0.7 diff |
| Number shared | 24 | 58 | 23 | 5 | 0 |
| Overlap of Lositan Loci with other filtering methods (total #loci = 256) |  |  |  |  |  |
|  | BayEnv 1% | Flip-Flop | 0.5 diff | 0.6 diff | 0.7 diff |
| Number shared | 5 | 100 | 118 | 43 | 3 |

Pairwise comparisons of major allele frequencies in populations closer to the central portions of the invaded range (“center populations” closer to the point of introduction, like Florida and The Bahamas) and those closer to the moving edge of the invaded range (“edge populations”) were used to detect specific loci for which frequencies were greater in the core of the invaded range than closer to the edge. From the list of loci that had a difference of 0.5 or more between maximum and minimum allele frequency, 115 had greater allele frequencies in Florida than in the USV, 127 had greater allele frequencies in Florida than in Honduras, 106 had greater allele frequencies in the Bahamas than in the USV and 122 had a greater allele frequency in the Bahamas than in Honduras. Additional pairwise comparison results showing counts of loci that overlap with different filtering requirements, including the outlier analyses described below are presented in Table 2.

### Outlier loci

LOSITAN analyses identified 256 loci as possible targets of directional selection (having an F_ST_ outside the upper bound of the 95% confidence interval, with a correction for multiple tests). *BayEnv 2.0* generated Bayes factors for the 12,759 analyzed loci. Taking the top 1% of loci from each bin captured 120 loci considered to have high enough Bayes Factors to be considered correlated to the linear regression model generated by the program *BayEnv*; taking the top 5% captured 616 loci. The top 5% of loci identified by *BayEnv* were then compared to the list of F_ST_ outliers generated by *LOSITAN* and were also compared to lists generated by the locus-specific diversity analyses described above, including the loci with a change from major to minor allele, and those with large differences between their maximum and minimum frequencies (Table 3). Of the 615 loci in the top 5% of BayEnv analysis, 24 were also identified as outliers by *LOSITAN* analysis. Of these 24 loci, 7 were putatively identified by Blast2GO. Several of these loci were identified by GO terms as being membrane proteins or involved in membranes (Table 4). Site frequency spectra and F_IS_ for different subsets of alleles had different distributions (Figure SI.VI.2, Figure SI.VI.3).

**Table 3.** Blast2GO results for those loci that overlapped between the BayEnv top 5% and the Lositan outlier results. In the GO terms, C = cellular component, P = biological process, F = molecular function.

| Locus | BLAST ID | GO terms |
| --- | --- | --- |
| 48803 | “glutamate receptor NMDA 2B” | C:postsynaptic membrane; P:ion transmembrane transport; C:integral component of membrane; C:cell junction; P:ionotropic glutamate receptor signaling pathway; F:ionotropic glutamate receptor activity; F:extracellular-glutamate-gated ion channel activity |
| 15012 | “proto-oncogene tyrosine-kinase Src isoform X1” | F:ATP binding; P:peptidyl-tyrosine phosphorylation; F:non-membrane spanning protein tyrosine kinase activity; P:response to yeast |
| 80176 | “coiled-coil domain-containing KIAA1407 homolog” | No GO Terms |
| 20821 | “KN motif and ankyrin repeat domain-containing 4-like” | No GO Terms |
| 11751 | “membrane progesterin receptor beta-like” | C:integral component of membrane |
| 54375 | “CD209 antigen E isoform X2” | C:membrane |
| 75133 | “PREDICTED: uncharacterized protein LOC103354480” | No GO Terms |

Of the loci identified as outliers in both *BayEnv* and *LOSITAN* (Table 3), four that were associated with GO terms were more closely scrutinized. In a BLAST-n query of the National Center for Biotechnology Information (NCBI) nucleotide database, only three of those four could be more specifically identified. These were, putatively, a glutamate receptor (locus 48803), a progestin receptor (locus 11751), and a tyrosine kinase (locus 15012) (SI-7).

## DISCUSSION

This study contains population genomic data generated using RAD-seq for the invasive lionfish, *Pterois volitans* collected from sites throughout the Caribbean Sea. Using 12,759 loci, we observed geographic patterns correlating diversity with distance from the point of invasion despite a lack of spatial metapopluation genetic structure. The most important of these patterns is the decrease of observed heterozygosity with distance from the point of invasion, despite no such pattern in expected heterozygosity, indicating a relationship between distance from the point of invasion and increased levels of disequilibrium. There are many factors that can lead to genetic disequilibrium, including essentially any violation of the Hardy-Weinberg assumptions. Considering the spatial relationship of the difference between observed and expected heterozygosity, the patterns of disequilibrium are likely related to spatial processes like expansion-driven genetic drift or, as discussed below, other aspects of population dynamics during expansion. No geographic metapopulation genetic structure was observed in either a principal component analysis or *fastSTRUCTURE* analysis and only minor differences in F_ST_ values were observed across nine populations in the Caribbean Sea. While mitochondrial data were consistent with previous genetic investigations concluding that a strong initial bottleneck was followed by mixing and that Caribbean currents may have helped to produce low levels of population differentiation in lionfish (Butterfield *et al.* 2015); the RAD-sequencing results for population structure did not find evidence of a genetic break between sites previously designated as Atlantic sites (Florida and The Bahamas) and those in the Caribbean (the rest of our study sites). The PCA and F_ST_ analyses of RAD-seq data presented here, rather, indicate that there is no structure. It is possible, therefore, that evidence of structure among basins seen in mtDNA data is solely driven by the absence of certain mitochondrial alleles in certain ocean basins. While structure was not seen in PCA analyses, directionality index results did point to the possibility that certain locations are more or less connected to the rest of the invaded range. For example, results indicated that the Dominican Republic may be more isolated from the population in Florida than expected and that Honduras is more connected. These results can serve as the basis for hypotheses about how population structure may develop in Caribbean lionfish populations through time as the region moves closer to an equilibrium state.

Elevated F_IS_ values in populations further from Florida could indicate cryptic structure (*e.g.*, the Wahlund Effect, Hartl and Clark, 1997). Population densities closer to the edge of the invasion could be lower than those closer to the center of the invaded range, which could account for the observed signature of cryptic structure. These patterns in F_IS_ could be the result of smaller population size, which could be one force driving cryptic structure in the populations near the moving range boundary. While F_IS_ values are not specifically elevated in populations where fish were sampled from multiple reefs, it is also possible that reef patchiness in different locations, or other sources of habitat heterogeneity could contribute to differences in F_IS_. In Three-Spined Stickleback, elevated patterns of F_IS_ have been linked to cryptic structure in newly colonized freshwater populations (Catchen *et al.* 2013).

### BLAST IDs of outlier loci

Using over ten thousand loci, we identified sites in the genome that break with equilibrium expectations. We identified 24 loci that are likely undergoing selection or strong genetic drift during expansion, including seven that were identified by BLAST, some of which were identified as membrane proteins by *Blast2GO* analysis. The majority of the identified outliers were not identified by BLAST. Because the knowledge of gene identity of our RAD-tags can be only cursory, we are unable to specifically disentangle signals for beneficial or deleterious mutations or alleles; however, we are able to infer that any present signals of allele surfing are not present in enough loci to result in an overall signal in major allele frequencies averaged over the loci sequenced and compared among populations throughout the range. While beyond the scope of this work, future analysis of longer sequence reads from paired-end RAD-seq data could aid in better locating and identifying these loci in fish genomes (Bourgeois *et al.* 2013).

In analyzing selection outlier results, it is common practice to identify loci that are shared among multiple outlier identification software programs and consider them to be stronger candidates for selection than those found only by one program. This is done because it is widely acknowledged that each method has its own limitations and that biases in specific models or assumptions about selection may skew the data when only one program is used (Lotterhos & Whitlock 2014). However, the F_IS_ and site frequency spectra for loci identified by *LOSITAN* and *BayEnv* result in visibly different genetic patterns (SI-6). *BayEnv* and *LOSITAN* use different metrics to find loci of interest, which look for “outliers” under different sets of expectations. LOSITAN identifies loci that have different allele frequencies (detected through deviations in F_ST_) in certain populations (Antao *et al.* 2008). *BayEnv*, in contrast, is based on detecting alleles that match a regression model with a certain parameter (in this paper, that parameter is distance from Florida), so it is likely to detect alleles with certain clines in the parameter space (Coop *et al.* 2010; Günther & Coop 2013). While these results may not have specific biological relevance because they are in part driven by the nature of different outlier analyses, they serve as a reminder that finding loci that overlap among multiple outlier analyses may result in the loss of importance nuance about certain patterns that are demonstrated by different analysis methods.

Of the loci that were identified as outliers by both *BayEnv* and *LOSITAN* and further confirmed through more extensive BLAST analysis, three in particular stand out as being potentially important biologically for lionfish because of the functional roles of these proteins. These are (1) locus 48803, which is likely located in a glutamate receptor; (2) locus 11751, which is potentially part of a membrane progestin receptor sequence; and (3) locus 15012, which was identified as being located in a proto-oncogene tyrosine kinase. These three loci could potentially be involved, respectively, in (1) learning and memory (Riedel *et al.* 2003); (2) gamete maturation (Tubbs *et al.* 2010), oocyte maturation (Zhu *et al.* 2013), and sperm hypermotility (Tan *et al.* 2014); and (3) cell division and growth (Newsted & Giesy 2000; Vivanco & Sawyers 2002). These results present a starting point for further research into the role of these loci in lionfish biology and gene function. Further research into regions of the genome under selection in the lionfish’s native range would be necessary to make any conclusions about the role of these loci in invasive lionfish evolution and adaptation.

### Study design affects genetic signals

The year of recruitment of individual fish is likely to affect genetic outcomes. For example, the fish sampled from the Cayman Islands—while collected in 2013—putatively recruited to the reef as early as 2005/2006, which would make them the oldest fish in the study (SI-1), which is interesting in light of the fact that observed heterozygosity of the Cayman Islands is lower than expected with the regression. This finding could be a result of the fact that the sampled Cayman fish are from an older age bracket, potentially representing a genetic cohort from earlier in the invasion. Fish sampled earlier could have lower diversity because they had just experienced the founder event characteristic of colonizing new locations and that population may quickly become more diverse after receiving more recruits from locations further behind the advance invasion wave. Therefore, the age and recruitment date of samples in population genetic studies of range expansion could be important for understanding the dynamics of invasion genetics. Population genetic papers usually assume a sufficiently long time scale of genetic change that the specific age class of individuals sampled is unimportant to the genetic conclusions (Bors *et al.* 2012); however, in rapid range expansions when genetic change is expected, differences of one or two years could change the expected genetic signals. Caution, therefore, may be necessary when drawing conclusions from datasets that include samples from many time points throughout an invasion, a practice that common in several recent lionfish mitochondrial DNA population genetic analyses (*e.g.,* Butterfield *et al.* 2015; Johnson *et al*. 2016).

This study is the first to employ RAD-seq to describe range expansion genetics in lionfish. While the development of next generation sequencing has accelerated the generation and analysis of large amounts of reduced representation genomic data, and the subsequent resolution of questions in the field of non-model species genomics (Reitzel *et al.* 2013; Therkildsen *et al.* 2013; Merz *et al.* 2013), analytical limitations due to the lack of whole genome sequences remain. For example, while the use of over 12,000 loci helped facilitate the identification of groups of loci that may be undergoing certain range expansion specific processes in this paper, it is possible that detecting specific clear examples of allele surfing (a rare allele becoming more common near a moving range boundary) may be difficult if only 0.31% of the genome is being sequenced (especially if, say, less than 1% of loci in the genome were demonstrating the pattern). So it is possible that allele surfing is still taking place and remain undetected by our analyses. The reduced representation strategy in this study samples less than one percent of the lionfish genome. Therefore, the likelihood of capturing loci that are experiencing allele surfing—unless there are many such loci—is low.

### Conclusions and implications

Range expansions, while an undeniably important force in shaping genetic diversity across the planet, have limited signatures in some species due to the specific context of the expansion. Here we have demonstrated that while not all the predicted patterns of expansion manifest themselves in the lionfish populations sampled, the range expansion process has led to disequilibrium closer to the range front. Populations of lionfish sampled in this study are well mixed and dispersal among sites is high, potentially precluding the detection of predicted decreases in allele frequency along the expansion axis. Genome-level analyses revealed low to no spatially explicit metapopulation genetic structure, yet genetic diversity throughout the invaded range was not homogeneous. While patterns of genomic diversity correlated with invasion pathway, observed heterozygosity decreased with distance from Florida while expected heterozygosity remained mostly constant, indicating population genetic disequilibrium correlated with distance from the point of invasion.

Ultimately, the lack of decreases in major allele frequency or allelic richness across the invaded range suggests that the process of expansion is unlikely to cause long-lasting limits to the adaptive potential of lionfish in their invaded range. It could also be inferred that signals of disequilibrium dissipate over time and space for the lionfish. Temporal comparisons of genetic diversity in a spatial context will be necessary to fully understand how a rapid invasion like that of the lionfish affects adaptive potential and the evolution of the species.

## ACKNOWLEDGEMENTS

We thank the many participants of the Gulf and Caribbean Fisheries Institute for providing lionfish samples from around the Caribbean region, as well as Dr. Bernard Castillo at the University of the Virgin Islands and Kristian Rogers at the Biscayne Bay National Park. We would like to acknowledge Alex Bogdanoff at NOAA, Beaufort NC, for assistance with sample acquisition; Camrin Braun at WHOI, for assistance with the calculation of oceanic distances between sites; Dr. Tom Schultz at Duke Marine Lab and Dr. Margaret Hunter at USGS for discussions concerning ongoing population genetic projects; and Jack Cook at the WHOI Graphics department for his assistance in generating maps of the study area. We would like to extend a special thank you to Dr. John Wakeley of Harvard University for assistance in the interpretation of data. This material is based upon work supported by the National Science Foundation Graduate Research Fellowship under Grant No. 1122374. Sequencing funding was provided in part by the PADI Foundation Grant No. 14904. Additional research support was provided by the Woods Hole Oceanographic Institution (WHOI) Ocean Ventures Fund, the Coastal Ocean Institute at WHOI, the National Science Foundation (OCE-1131620 to TMS), and the James Education Fund for Ocean Exploration within the Ocean Exploration Institute at WHOI.

